# Single-cell time-resolved multi-omics reveal apoptotic and ferroptotic heterogeneity during foam cell formation

**DOI:** 10.1101/2023.03.30.534700

**Authors:** Yiwen Wang, Ling Lin, Liang Qiao

**Affiliations:** Department of Chemistry, Shanghai Stomatological Hospital, and Zhongshan Hospital, Fudan University, Shanghai 200000

**Keywords:** Single-cell mass spectrometry, Macrophage, Atherosclerosis

## Abstract

Macrophage-derived foam cell plays a pivotal role in the plaque formation and rupture during the progression of atherosclerosis. Foam cells are destined to divergent cell fate and functions in response to external stimuli based on their internal states, which however is hidden in the traditional studies based on population of cells. Herein, we used time-resolved and single-cell multi-omics to investigate the macrophage heterogeneity along foam cell formation. Dynamic metabolome and lipidome outlined the dual regulating axis of inflammation and ferroptosis. Single cell metabolomics and lipidomics further demonstrated a macrophage continuum featuring a differed susceptibility to apoptosis and ferroptosis. Using single-cell transcriptomic profiling, we verified the divergent cell fate toward apoptosis or ferroptosis. Therefore, the molecular choreography underlying the divergent cell fate during foam cell formation was revealed, which is of high significance for the understanding of the pathogenesis of atherosclerosis and development of new drug targets.

## Introduction

Atherosclerosis is a chronic inflammatory disease driven by the maladaptive lipid metabolism and inflammatory response of macrophages^1,2^. At the initiation of atherosclerotic lesions, circulating monocytes differentiate into adherent macrophages; as the lesion expands, macrophages take in lipids to form foam cells, which leads to the ultimate plaque and thrombus in the artery^3,4^. Foam cell death plays a pivotal role in the plaque formation and rupture. According to the classical view, lesional macrophages were destined to apoptosis, leading to secondary necrosis and the ultimate plaque formation^1,3^. Further studies have suggested a potential necroptosis, autophagy, pyroptosis and ferroptosis in lesional macrophages^5–8^. The programmed cell death of foam cells has been rated as pro-atherogenic (apoptosis in advanced lesions, pyroptosis, necroptosis and ferroptosis) or anti-atherogenic (apoptosis in early lesions and autophagy)^9^. Nevertheless, current studies based on population of cells hide the molecular choreography that governs the divergent cell fate^10,11^.

Investigations with lineage tracing and time-lapse fluorescence imaging have demonstrated the single-cell heterogeneity in apoptosis susceptibility regulated by p53 dynamics^12–14^. Using the time-lapse intravital microscopy, Robbins et al. mapped the development of monocytes from extramedullary niches and into foam cells in the murine model of atherosclerosis^15^. While time-lapse imaging enables continuous long-term single-cell quantification for understanding the kinetics of molecules of interest, emerging cell snapshot techniques such as single-cell RNA sequencing and single-cell mass spectrometry provide omics readout with much higher dimension of molecular information. Investigations with fate mapping and single-cell RNA sequencing have demonstrated the developmental patterning and the transcriptional dysregulation of monocytes in the progression and regression of atherosclerosis^16,17^. Single-cell RNA sequencing, combined with upstream single-cell assay for transposase-accessible chromatin sequencing (scATAC-seq) and downstream cellular indexing of transcriptomes and epitopes by sequencing (CITE-Seq), has defined foam cell subtypes based on the integrated dimensions of epigenetics, transcripts and epitopes^18–20^. While single-cell sequencing revealed various macrophage subtypes with predicted functions, its combination with downstream single-cell proteomic, metabolomic and lipidomic profiling would reflect in a more direct and dynamic way the molecular mechanism underlying the heterogeneity during foam cell formation.

To date, current dynamic metabolomic and lipidomic studies on lesional macrophages were mainly carried out in bulk. Macrophages work as a lipid scavenger and immune regulator by dynamic lipid and metabolic reprogramming in response to the microenvironment during the progression of atherosclerosis^21,22^. Metabolic flux and dynamic lipidomic analysis have helped unveil the reprogramming of immunometabolism and lipid metabolism in macrophages. Using metabolic flux analysis, Haschemi et al. demonstrated an accelerated glycolysis flux toward pentose phosphate pathway (PPP) in lipopolysaccharide-activated macrophages^23^. The dysregulation of multiple amino acid anaplerosis pathways have been further proposed for macrophage polarization^24,25^. Using metabolic flux analysis, Argus et al. demonstrated distinct reprogramming of the *de novo* biosynthesis and elongation of fatty acids in macrophages treated with different pro-inflammatory stimuli^26^. Using a time-resolved global lipidomic profiling, Dennis et al. demonstrated a cascade starting from eicosanoids toward sterols, sphingolipids, glycerophospholipids and glycerolipids in macrophages activated by pro-inflammatory stimuli^27^.

Herein, we used time-resolved and single-cell multi-omics profiling to investigate the metabolic and lipid reprogramming and to depict macrophage heterogeneity on molecular level (Fig. 1). Dynamic metabolome and lipidome outlined the dysregulation in the immunometabolism of PPP, TCA and amino acid anaplerosis and metabolism of ether phospholipids. Single-cell metabolite and lipid profiling depicted a macrophage continuum featuring a differed susceptibility to apoptosis and ferroptosis. Single-cell transcriptomic profiling verified the divergent cell fate toward apoptosis and ferroptosis in the late-stage foam cells.

**Figure 1.**
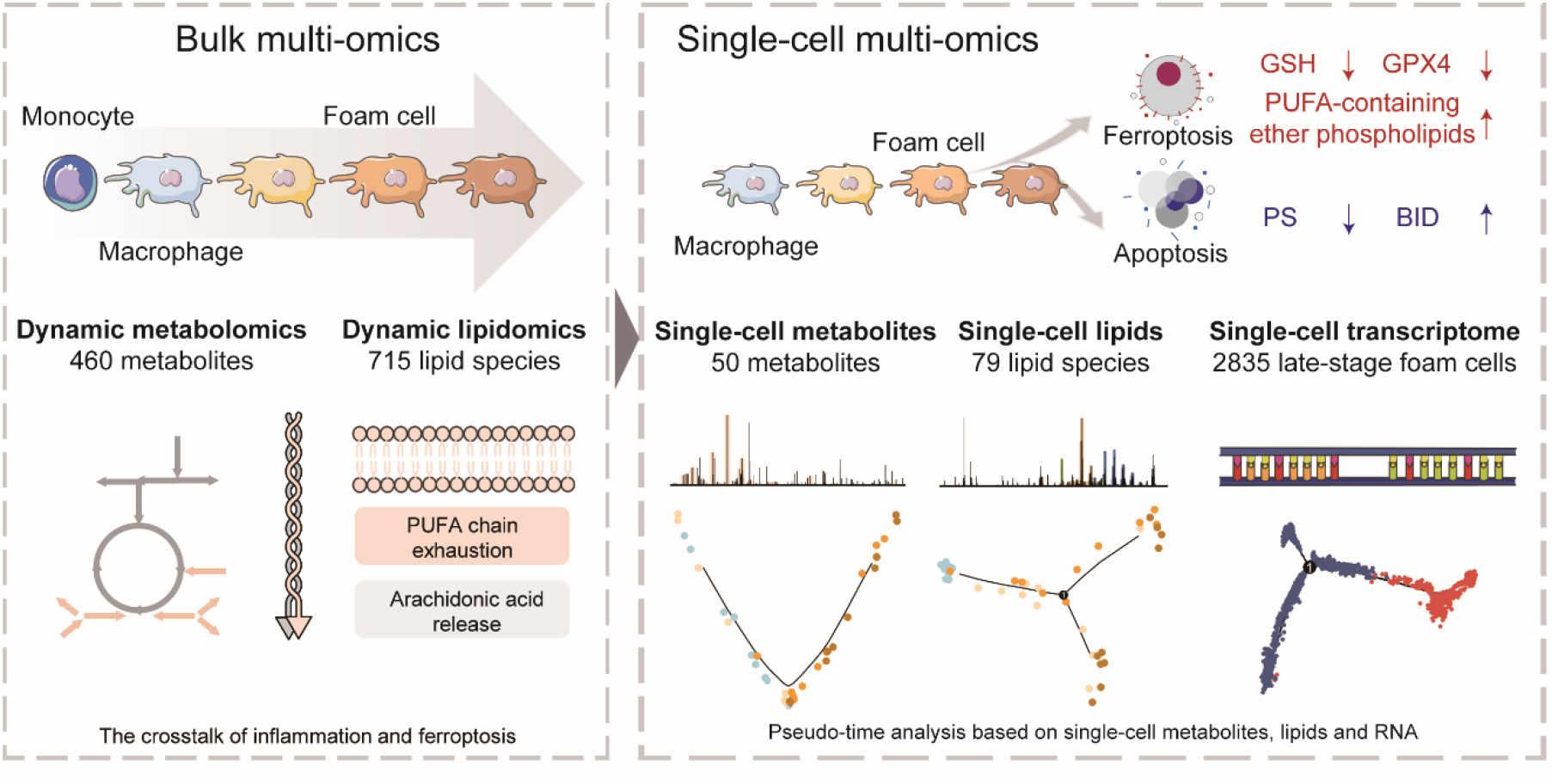
Overview of the study design. Time-resolved metabolomics and lipidomics are firstly applied on population of cells during the foam cell formation, revealing the dysregulation in the immunometabolism of PPP, TCA and amino acid anaplerosis and metabolism of ether phospholipids. Single cell time-resolved metabolomics and lipidomics are further adopted, indicating the differed susceptibility of foam cells to apoptosis and ferroptosis. Single cell transcriptomics verified the divergent cell fate toward apoptosis and ferroptosis in the late-stage foam cells.

## Results

### Metabolomic and lipidomic cascades in monocyte differentiation and foam cell formation

In this study, THP-1 cell line was used to prepare macrophages and foam cells. We used the THP-1 cell line to mimic the monocytes (0T), differentiated the monocytes into the macrophages (1C) and then incubated the macrophages with oxidized low-density lipoprotein (oxLDL) for different hours to generate the foam cells of early, middle and late stage (2E, 3M and 4L, respectively) (Fig. 1 and *SI appendix*, Fig. S1). Untargeted metabolomic and lipidomic profiling were performed on the five groups of cells (0T, 1C, 2E, 3M and 4L) with three biological replicates each, identifying 460 metabolites, and 715 lipid species with 290 known molecular species. The identified metabolites were then subjected to superclass and pathway enrichment by KEGG. The metabolites were enriched in superclasses of benzoids, organic acids, nucleic acids, fatty acyls, etc. (Fig. S2*A*). The metabolic pathways were enriched in the purine metabolism, alanine, aspartate and glutamate metabolism, pyrimidine metabolism, etc. (*SI appendix*, Fig. S2*B*). The lipidomic dataset covers 24 subclasses including neutral lipids, glycerophospholipids, sphingolipids and fatty acids (*SI appendix*, Table S1).

PLSDA analysis on metabolite and lipid species level has demonstrated a sound separation among monocytes, macrophages and foam cells of three stages (Fig. 2*A*). According to the ANOVA test, a total of 159 (35% of all the identified) metabolites and 340 (48%) lipid species demonstrated significant changes throughout the differentiation and foam cell formation compared to the macrophages, suggesting a large-scale metabolic and lipid rewiring (Fig. 2*B*). More metabolites and lipids were upregulated during the monocyte differentiation to macrophage. During the foam cell formation from macrophage, a dominating upregulation in metabolites and lipid species was observed. On the subclass level of lipids, an upregulation in sphingolipids (e.g. LacCer, GlcCer) and anionic glycerophospholipids (PS and PI) was observed during the differentiation of monocyte to macrophage, while a decrease in sphingolipids (LacCer, GlcCer) and an increase in neutral lipids (ST, TG and DG) occurred throughout the foam cell formation (Fig. 2*C*). Notably, the increase in sterols and triglycerides resonates with the prior knowledge of macrophage’s lipid loading under hyperlipidemia^28^.

**Figure 2.**
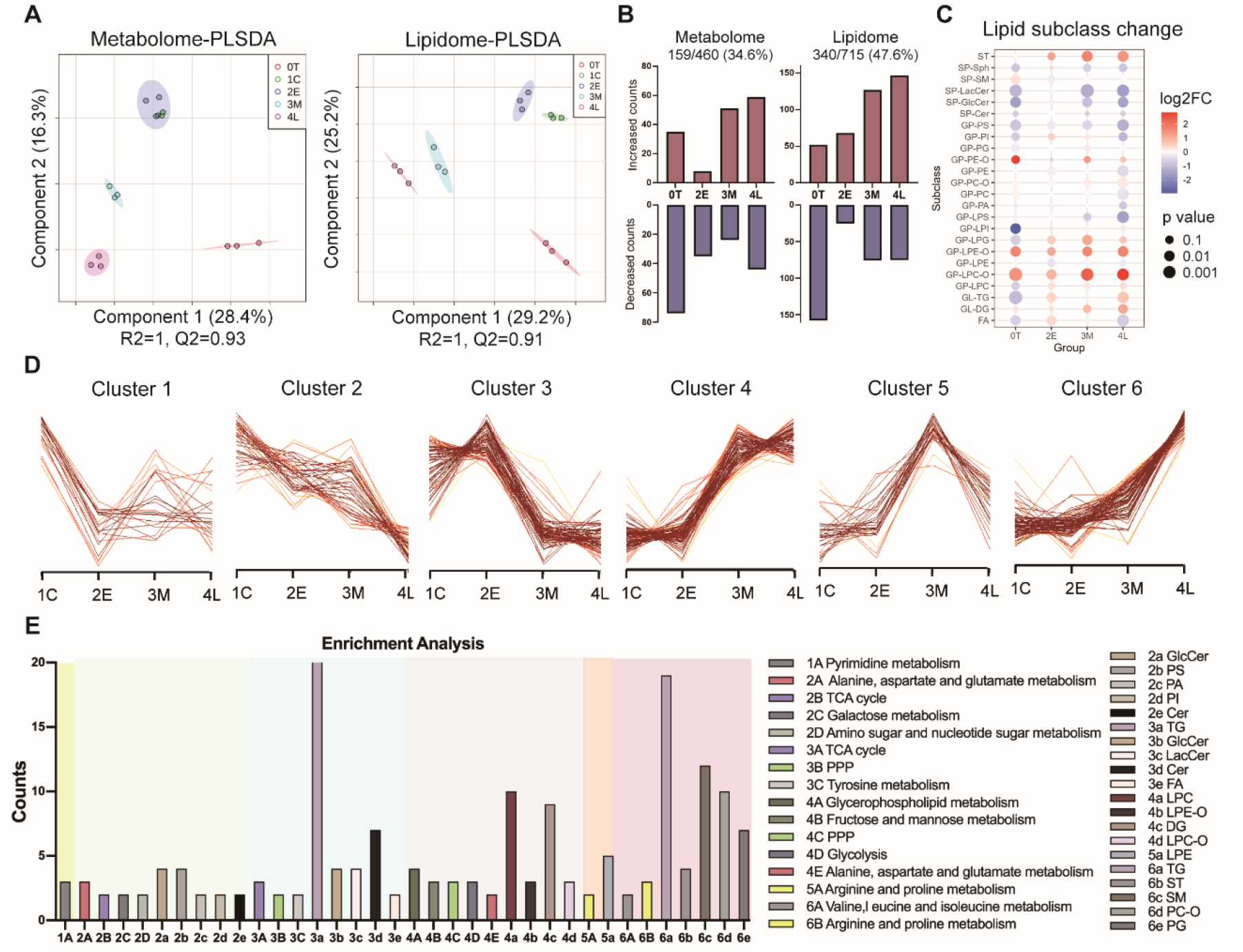
Metabolomic and lipidomic changes throughout the differentiation of monocyte and foam cell formation: (*A*) PLSDA score plots based on metabolites and lipid species. (*B*) Dynamic changes on metabolite and lipid species level. 0T, 1C, 2E, 3M and 4L represent monocytes, macrophages and foam cells of early, middle and late stages, respectively. 0T, 2E, 3M and 4L were compared with the 1C group, respectively. (*C*) Dynamic changes on lipid subclass level. Fold change was calculated against the 1C group. (*D*) Clustering of the differential metabolites and lipids. (*E*) Metabolic pathway and lipid sub-classes enrichment analysis of the clusters in (*D*). 1A represents metabolites in cluster 1, while 1a represents lipids in cluster 1. The others follow the same rule. PPP, pentose phosphate pathway.

We further focused on the metabolic and lipid reprogramming during foam cell formation. Differential metabolites and lipid species were then clustered by their abundance change along foam cell formation. Six molecular clusters were identified (Fig. 2*D*), delineating miscellaneous biological processes encompassing endocytosis, apoptosis, inflammation and ferroptosis (Fig. 2*E*). Among them, Cluster 1 (n = 21) features an early depression; Cluster 2 (n = 43) displayed a maintained downregulation; Cluster 3-6 (n = 68, 85, 30 and 89, respectively) demonstrated a delayed response (Fig. 2*D*). Cluster 1 was enriched in molecules associated with pyrimidine metabolism (Fig. 2*E*), which is recognized as a property of M2 type^29^. For Cluster 2, the dysregulation of anionic phospholipids (PS, PA and PI) reflected the perturbation in membrane charge density and curvature, which are related to endocytosis and cholesterol efflux^30,31^. PS(18:0/18:1), the molecular species identified to interact with cholesterol to tune the transbilayer distribution of lipids, displayed a maintained decrease^32^ (Fig. 2*D* and *SI appendix*, Fig. S3*A*). The exhaustion of PS species suggested the caspase-mediated apoptosis in response to long-term oxLDL exposure^33^. The sustained depression of nucleotide sugar metabolism (Cluster 2), on the other hand, reflected the suppressed N-glycosylation (*SI appendix*, Fig. S3*B*), which is established as an anti-inflammatory M2 marker^24,34^. For Cluster 4, an increase in lipid metabolism reflected the response to increasing lipid import for striking a homeostasis. For Cluster 6, the delayed rise in branched chain amino acid (BCAA) pathway suggested a compensatory response to generate anti-atherogenic leucine that can lower TG content by blocking very low density lipoprotein (VLDL) endocytosis and TG biosynthesis and promoting cholesterol efflux.^35^

The clustering analysis observed either a stepwise or a divergent dysregulation in pathways of pentose phosphate pathway (PPP, Cluster 3 and 4), TCA cycle (Cluster 2 and 3), arginine and proline metabolism (Cluster 5 and 6), alanine, aspartate and glutamate metabolism (Cluster 2, 3 and 4) and ether lipid metabolism (Cluster 4, 5 and 6). We then delved into the dysregulation of the metabolites and lipid species involved in the aforementioned pathways.

### Inflammation and ferroptosis along the foam cell formation

For the immunometabolism, a channeling PPP, broken TCA cycle and defective amino acid anaplerosis were observed (Fig. 3*A*). The downregulation of sedoheptulose-7P indicated the suppressed carbohydrate kinase-like protein (CARKL), which is required for refocusing on the glycolysis and M1-like metabolic reprogramming^36^. The elevated branch of PPP (from glucono-1,5-lactone to gluconate) fueled the NADPH production and downstream ROS generation. TCA witnessed a stepwise downregulation with the second carbon oxidation (from fumarate to malate) decreased at the early stage of foam cell, and the upstream pyruvate oxidation and first carbon oxidation (from pyruvate to oxo-glutarate) suppressed at the middle stage of foam cell. The dysregulation of the intertwined glycolysis, PPP and TCA suggested an ever-increasing inflammatory M1 phenotype and a diminishing M2 type since the early stage of foam cell, in accordance with the prior recognition that oxLDL-induced foam cells acquire a M1-like phenotype^37^.

**Figure 3.**
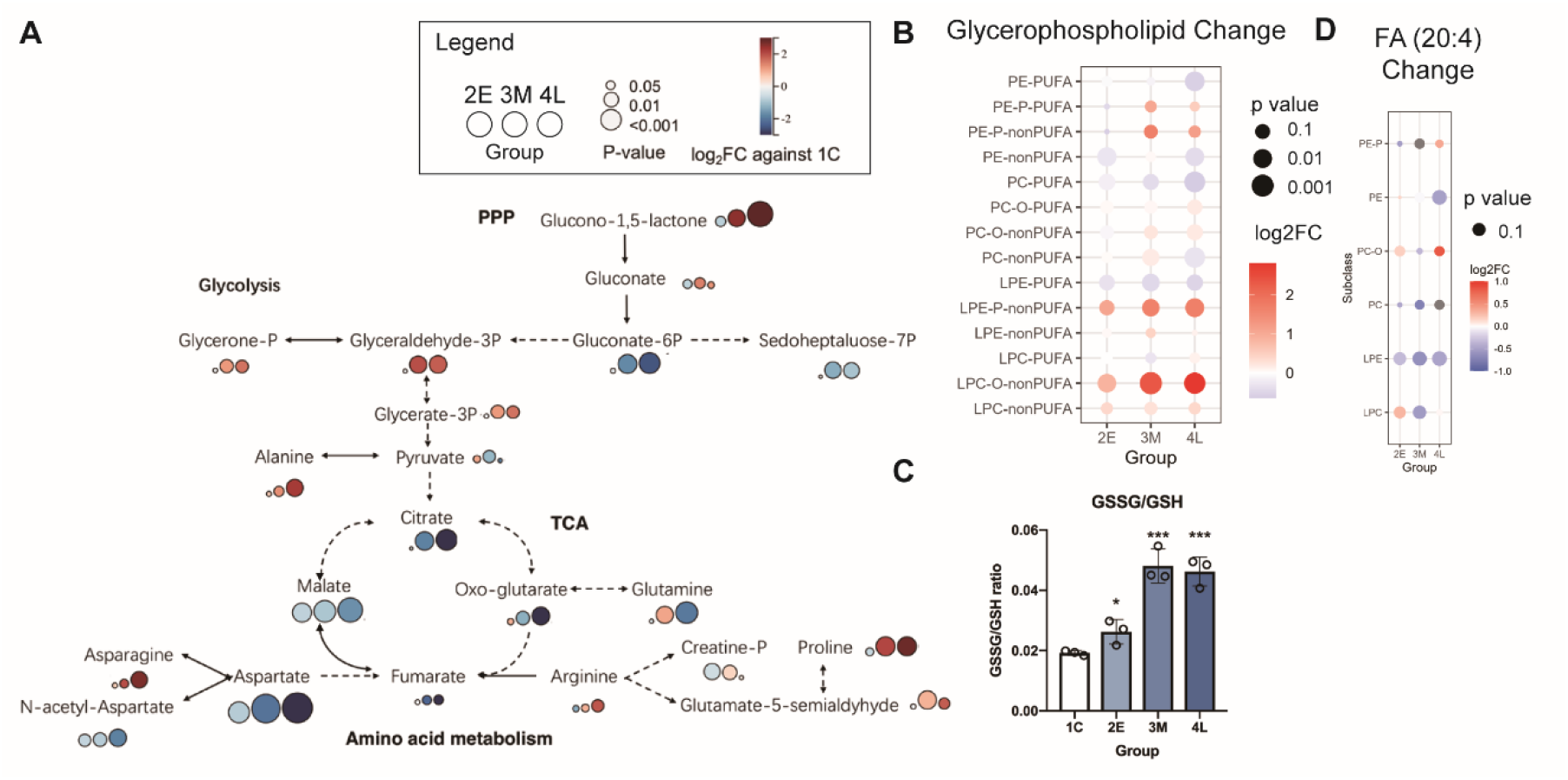
Inflammation and ferroptosis during foam cell formation: (*A*) Regulation of metabolites involved in the immunometabolism of glycolysis, PPP, TCA and amino acid metabolism. The full lines and dotted lines represent one-step and multiple-step reactions, respectively. The color and size of the bubbles represent the fold change and significance against 1C group, respectively. The circles from left to right represent early, middle and late stage of foam cells, respectively. (*B*) Dynamic changes of glycerophospholipids. Fold change and significance were calculated against the 1C group. (*C*) GSSG/GSH levels in macrophages and foam cells of three stages. Error bars represent the standard deviation. Significance was calculated against the 1C group with “*” and “***” in represent of p-values less than 0.05 and 0.001, respectively. (*D*) Dynamic changes in glycerophospholipids with a FA (20:4) chain. Fold change and significance were calculated against the 1C group.

In amino acid metabolism, the defective aspartate anaplerosis led to the accumulation of asparagine, which promoted the interleukin-1 *β* (IL-1 *β*) secretion for macrophage to acquire a pro-inflammatory phenotype^38^ and the compensatory regulation of atheroprotective alanine^35^. The dysregulated arginine and proline metabolism pathway featured a stable creatine shunt and an increasing proline biosynthesis. Arginine metabolism represents a classical signature for macrophage polarization, fueling either the production of reactive nitrogen species via inducible nitric oxide synthase (iNOS) in pro-inflammatory M1 type or feeding downstream proline and polyamine synthesis via arginase1 (ARG1) in anti-inflammatory M2 type^39^. The increasing flux siphoned from arginine to proline (*SI appendix*, Fig. S4*A*) thus suggested a relatively suppressed iNOS, a synthase reported to orchestrate the TCA dysregulation, glutaminolysis^40^ and ferroptosis susceptibility^41^. Glutaminolysis experienced a gradual decrease till the middle stage and an insignificant rebounce in the late stage (*SI appendix*, Fig. S4*B*). Glutaminolysis represents the most upregulated pathway in M2 type, modulating ferroptosis^42^ and efferocytosis^25^. The dysregulated glutaminolysis therefore reflected an impaired efferocytosis function and an upregulated ferroptosis susceptibility at late stage. The defective amino acid anaplerosis suggested additional regulation axes along the foam cell formation apart from the well-recognized polarization toward the classical M1 type.

For the ether lipid metabolism, the dysregulation rippled from lysoplasmalogens without polyunsaturated fatty acid (PUFA) chains toward the following diacyl phospholipids with PUFA chains. (Fig. 3*B*). The rise in non-PUFA LPE-P and LPC-O reflected an increasing hydrolysis to release PUFA. A downregulation in lipids with PUFA chain and an upregulation in ether-linked lipids suggest the response to oxidative stress. The increase of GSSG/GSH ratio reflected a suppression in Glutathione Peroxidase 4 (GPX4) and an increase in ferroptosis along the foam cell formation (Fig. 3*C*). The release of pro-inflammatory arachidonic acid mainly occurred from PE and LPE species at the late stage (Fig. 3*D*), suggesting the crosstalk between pro-inflammatory immunometabolism and ether lipid metabolism^43^.

Using the dynamic metabolomic and lipidomic profiling, we monitored the metabolic and lipid reprogramming throughout the monocyte differentiation and foam cell formation, focused on the dysregulation of immunometabolism and ether lipid metabolism along foam cell formation, and dissected the interplay between the dual axes of inflammation and ferroptosis.

### Single-cell metabolite and lipid profiling reveal differed susceptibility to ferroptosis and apoptosis of foam cells

Given the inflammatory and ferroptotic regulation revealed in bulk, we further performed a single-cell time-resolved metabolite and lipid profiling to investigate the heterogeneity along foam cell formation. We performed the single-cell mass spectrometry analysis based on a micromanipulation stage^44^ and nano electrostatic spray ionization (nano-ESTASI) mass spectrometry^45^ (*SI appendix*, Fig. S5). Single-cell metabolite and lipid profiles (*SI appendix*, Fig. S6 and Fig. S7) were extracted from raw data and the identity of the metabolites and lipid species was attributed and validated by the metabolome and lipidome acquired in bulk based on the high-resolution molecular weight. Ten single cell replicates were included for each of the four groups (1C, 2E, 3M and 4L). Single cell metabolite and lipid profiling of macrophages and foam cells covered a total of 79 lipid species and 50 metabolites. Metabolic pathways, such as lipid metabolism and amino acid and nucleotide metabolism (*SI appendix*, Fig. S8*A*), and lipid subclasses, such as TG and PE-P species (*SI appendix*, Fig. S8*B*) were enriched in the single-cell metabolite and lipid datasets. Time-series clustering analysis on single-cell metabolite and lipid datasets observed a pattern reminiscent of the bulk analysis, featuring a constant rise in neutral lipids, a maintained decrease in anionic phospholipids and a divergent dysregulation in amino acid anaplerosis (*SI appendix*, Fig. S9).

UMAP and PLSDA analysis on single-cell metabolite and lipid species has both demonstrated a continuum among macrophages and foam cells of three stages (Fig. 4*A* and *SI appendix*, Fig. S10). For the single-cell lipid profiling, a gradual increase of TG species was observed along the foam cell formation, while certain ether lipids and PE species were specifically enriched in macrophages (Fig. 4*A* and *SI appendix*, Fig. S11). For the singe-cell metabolite profiling, a maintained increase of glycerophosphocholine (GPC) was observed along the foam cell formation. A gradual exhaustion of traumatic acid (TA), an antioxidant recognized to alleviate phospholipid peroxidation, as well as a decompensatory dysregulation of GSH were observed along the foam cell formation, reflecting an unresolved response to oxidative stress. Using the pseudo-time analysis, we reconstructed the developmental trajectory of macrophages from the homeostatic state to the foam-cell state, based on the single-cell metabolite and lipid features (Fig. 4*B*). The heatmap defined clusters of differential molecules based on their changes along the single-cell pseudo-time (Fig. 4*C*). TG species together with metabolites of lipids demonstrated a constant upregulation, reflecting an increasing lipid retention along the foam cell formation, while glycerophospholipids and glutathione showed a maintained downregulation, resonating with the exhaustion of anionic phospholipids in apoptosis and antioxidants in ferroptosis. The pseudo-time analysis on single-cell lipid species also delineated a split in foam cells of the late stage. The PLSDA analysis and volcano plot of the two subpopulations of the late-stage foam cells based on single-cell lipids suggested that ether phospholipids and TG species account for the split (*SI appendix*, Fig. S12). The divergent expression of ether phospholipids and heterogeneous expression of glutathione in the late-stage foam cells suggested that there might exist a heterogeneity in foam cell ferroptosis (Fig. 4*D*).

**Figure 4.**
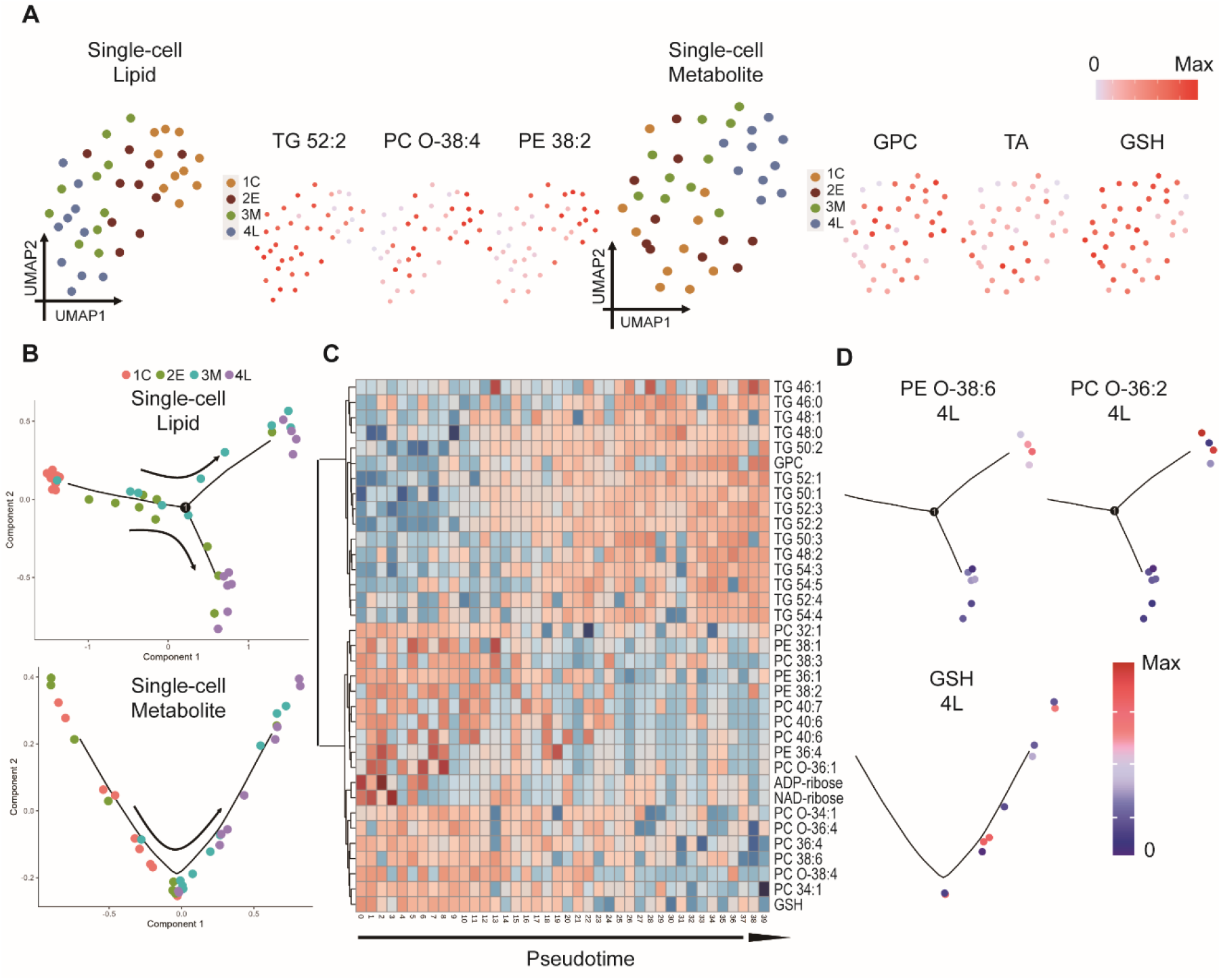
Single-macrophage continuum during the foam cell formation: (*A*) UMAP analysis based on single-cell lipid features and metabolite features. The UMAP layouts were colored by the cell group and then the abundance of lipid and metabolite features. (*B*) Pseudo-time trajectory of macrophages and foam cells based on single-cell lipid and metabolite profiling. Arrows indicate the direction of pseudo time increase. (*C*) Heatmap of differential lipid species and metabolites for all cells along the pseudotime. (*D*) Expression of certain lipid and metabolite features in foam cells of late stage.

We further focused on the non-continuous heterogeneity residing in the foam cell lipids and metabolites. The variability of lipid species and metabolites on single-cell level was summarized by their coefficients of variance (CV). Five molecular clusters have been defined according to their changes of CV across different stages of foam cells (Fig. 5*A*). Cluster 1 features a maintained low CV; Cluster 2 displayed an early decrease in CV; Cluster 3 demonstrated a high CV at the early stage; Cluster 4 showed peak CV in the middle stage; Cluster 5 displayed a delayed CV increase at the late stage. According to the enrichment analysis (Fig. 5*B*), PC and TG species of high abundance populated the Cluster 1 and 2. Amino acid and purine metabolism were enriched in the Cluster 2. Nicotinate and nicotinamide metabolism was enriched in Cluster 3. Membrane phospholipids such as PE, PS, PC, SM and ether lipids were enriched in Cluster 4 and 5. Amino acid metabolism, glutathione metabolism, purine and pyrimidine metabolism, TCA cycle and glycerophospholipid metabolism were enriched in Cluster 4 and 5. Among them, the enrichment of PE, PS and ether lipid subclasses and glutathione metabolism pathway again suggests high variability in foam cell apoptosis and ferroptosis at the middle and late stages.

**Figure 5.**
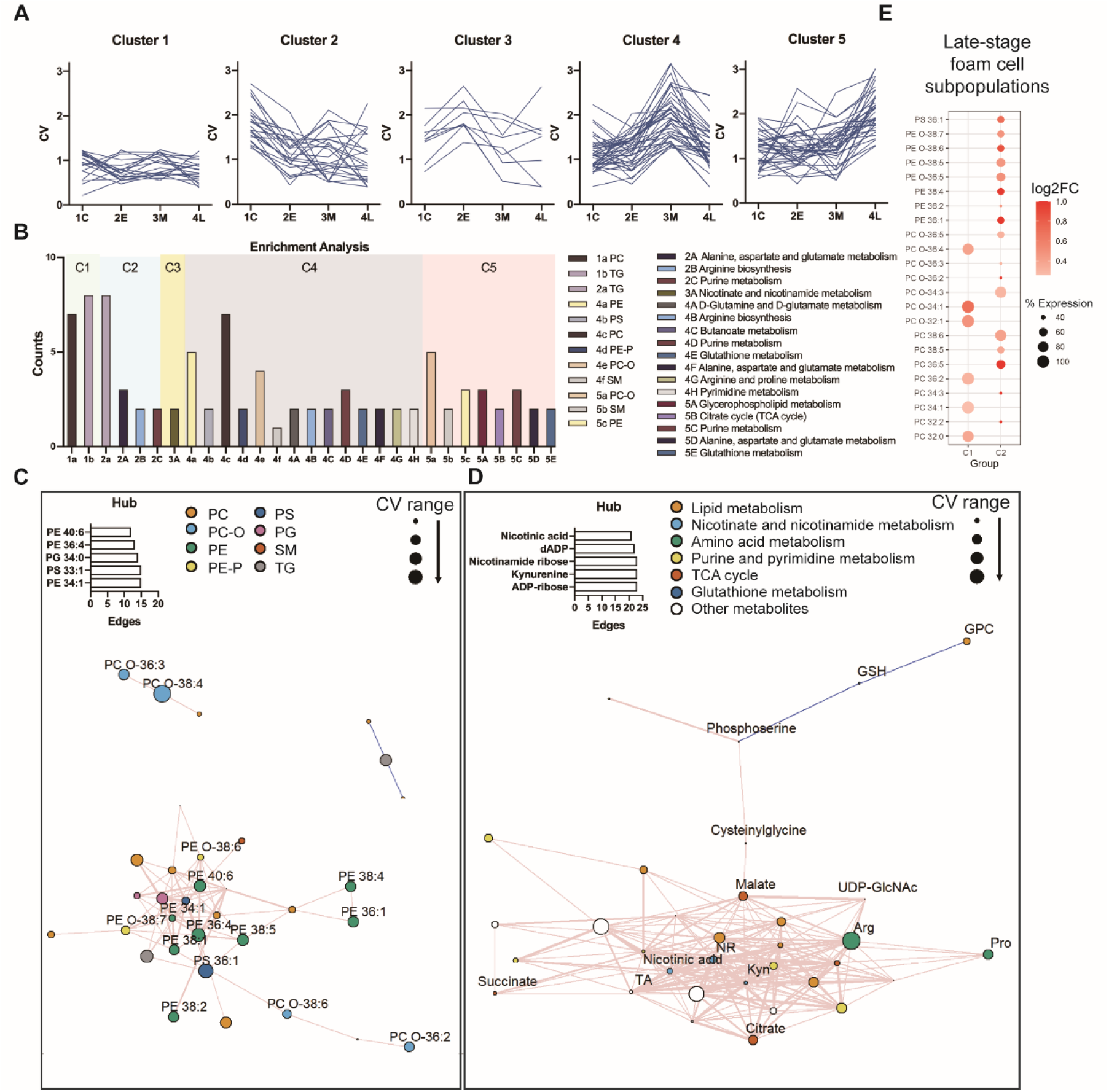
Single-macrophage heterogeneity during foam cell formation. (*A*) Clustering of single-cell features based on the change of coefficient of variance along the foam cell formation. (*B*) Metabolic pathway and lipid sub-classes enrichment analysis of the clusters in (*A*). (*C, D*) Lipid covariation network with lipid species labelled in color according to their subclasses and metabolite covariation network with metabolites labelled in color according to their pathways. The size of vertices represents the CV range (peak CV minus 1C CV) of the molecules. Edges represent spearman correlation with a p-value < 0.05. Edges of positive and negative correlation were labelled in rosy and blue color, respectively. The width of edges indicated the coefficients of correlation. The barplot indicates the edge count of the top 5 hubs in the network. (*E*) Lipid expression dotplots. UDP-GlcNAc, uridine diphosphate N-acetylglucosamine; NR, nicotinamide ribose; Arg, arginine; Pro, proline; Kyn, kynurenine; TA, traumatic acid.

To test whether the metabolic pathways and lipid subclasses are coordinately modulated, we performed a correlation analysis on the metabolites and lipid species of high CV in foam cells (cluster 3, 4, 5). The correlation analysis showed a lipid coregulation of PE, PS and ether phospholipid species and a metabolite coregulation centered with nicotinate and nicotinamide metabolism along the foam cell formation (Fig. 5*C* and 5*D*). PE and PS species populated the hubs in the lipid covariance network, extending to other membrane phospholipids as PC, SM and ether lipids. PE and PS are metabolically related aminophospholipids^46^ and work synergistically in tuning membrane curvature^31^ and binding to signaling proteins^47^. Therefore, the coregulation of PE and PS species suggested heterogeneity in the apoptotic and efferocytotic state of foam cells. The covariance of PE and PS species extended to ether phospholipids. PE-P and PC-O species, with their labile ether bond, are susceptible to lipid peroxidation^48,49^. The coregulation of ether phospholipids therefore suggested heterogeneous ferroptosis in foam cells aside from apoptosis. In the metabolite covariance network, arginine and proline demonstrated apparent CV changes during foam cell formation. The coregulation of arginine and proline suggested a detour from iNOS dependent nitric oxide (NO) synthesis toward ARG1 based proline biosynthesis. iNOS and NO have been recognized to render macrophages resistance to ferroptosis^41^. The covariance of arginine and downstream proline therefore suggested heterogeneity in inflammation and ferroptosis susceptibility. Nicotinate and nicotinamide metabolism populated the hubs in the metabolite covariance network, connecting amino acid metabolism, TCA cycle, purine metabolism and lipid metabolism. Nicotinamide adenine dinucleotide (NAD^+^), a kernel cofactor in glycolysis and oxidative phosphorylation, is derived from the *de novo* pathway via kynurenine and nicotinate, and the salvage pathway via nicotinamide ribose. Hence, the interlaced hubs of nicotinate, nicotinamide ribose and kynurenine together with their extensions resonated with the prior knowledge that NAD^+^ metabolism plays a potential regulatory role in inflammation, phagocytosis and respiration^50^.

Based on the coregulation of functional lipid species and metabolites, we tentatively divided macrophages into subpopulations annotated with different biological processes. Focused on the heterogeneous lipid expression in foam cells of the late stage, we witnessed a subpopulation displaying exhaustion of PE and PS species (C1) and a subpopulation enriched in ether phospholipids with PUFA chain (C2) (Fig. 5*E*), suggesting a divergence in cell fate toward apoptosis and ferroptosis.

### Single-cell transcriptomics verifies divergence in apoptosis and ferroptosis of foam cells

We performed a single-cell transcriptomic sequencing to verify the divergent cell fate in a high-throughput manner. A total of 2835 single late-stage foam cells were recovered after the quality control. UMAP analysis demonstrated two cell clusters with distinctive expression of ferroptosis and apoptosis marker genes (Fig. 6*A* and 6*C*). Cluster 0 featured an overexpression of apoptotic genes (e.g., BAD, BCL2L11) and ferroptosis suppressor genes (e.g., FTH1, GPX4); Cluster 1 featured an overexpression of ferroptosis driver genes (e.g., TFRC) (Fig. 6*B*). To identify cluster-specific gene expression, we checked the enrichment of gene sets at the single-cell level by calculating gene expression scores (*SI appendix*, Fig. S13). Several pathways overlaid with the defined cell clusters: Cluster 0 overexpressed genes involved with cell response to redox, peroxisome, hypoxia, electron transfer chain (ETC) in mitochondria, iron transport and uptake, and M1 type; Cluster 1 overexpressed genes implicated in acute inflammatory response, regulation of cholesterol efflux and inorganic ion import across plasma membrane. Further, we compare our foam cell subpopulations with macrophage subpopulations from recent publications on single-cell transcriptomic analysis of human atherosclerotic plaques^20,51^ (*SI appendix*, Fig. S14). The apoptosis-like cluster 0 in our dataset showed significant overlap with the C2 cluster in Fernandez et al.’s clinical dataset^20^, which was enriched in plaques from symptomatic patients and related to the iron clearance in the areas of intraplaque hemorrhage, the C5 cluster, which was enriched in plaques from asymptomatic patients and annotated as the foamy macrophages and the C9, 10 cluster, which were enriched in plaques from symptomatic patients and related to M2 phenotype (*SI appendix*, Fig. S14*A*). Cluster 0 also showed significant overlap with the My 0 cluster, which was annotated as recently recruited macrophages, and My 2 cluster, which was annotated as foamy macrophages, in Depuydt et al.’s publication^51^ (*SI appendix*, Fig. S14*B*). The ferroptosis-like cluster 1, on the other hand, showed significant overlap with the My 1 cluster annotated as pro-inflammatory macrophages and My 2 cluster. The overlap demonstrated a decent concordance between our *in vitro* model and human patients. To further confirm whether ferroptosis takes place in human plaque macrophages, we focus on the expression of ferroptosis driver and suppressor genes in the clusters showing significant overlaps (*SI appendix*, Fig. S14*C* and S14*D*). C2 cluster hit with the ferroptosis suppressor gene FTH1. My 2 cluster showed sound overlap with ferroptosis suppressor genes (FTH1, GPX4 and HSPB1), while My 0 and My 1 cluster hit with ferroptosis driver genes. The overlap with ferroptosis marker genes confirmed that the heterogeneity of ferroptosis exists in human plaque macrophages.

**Figure 6.**
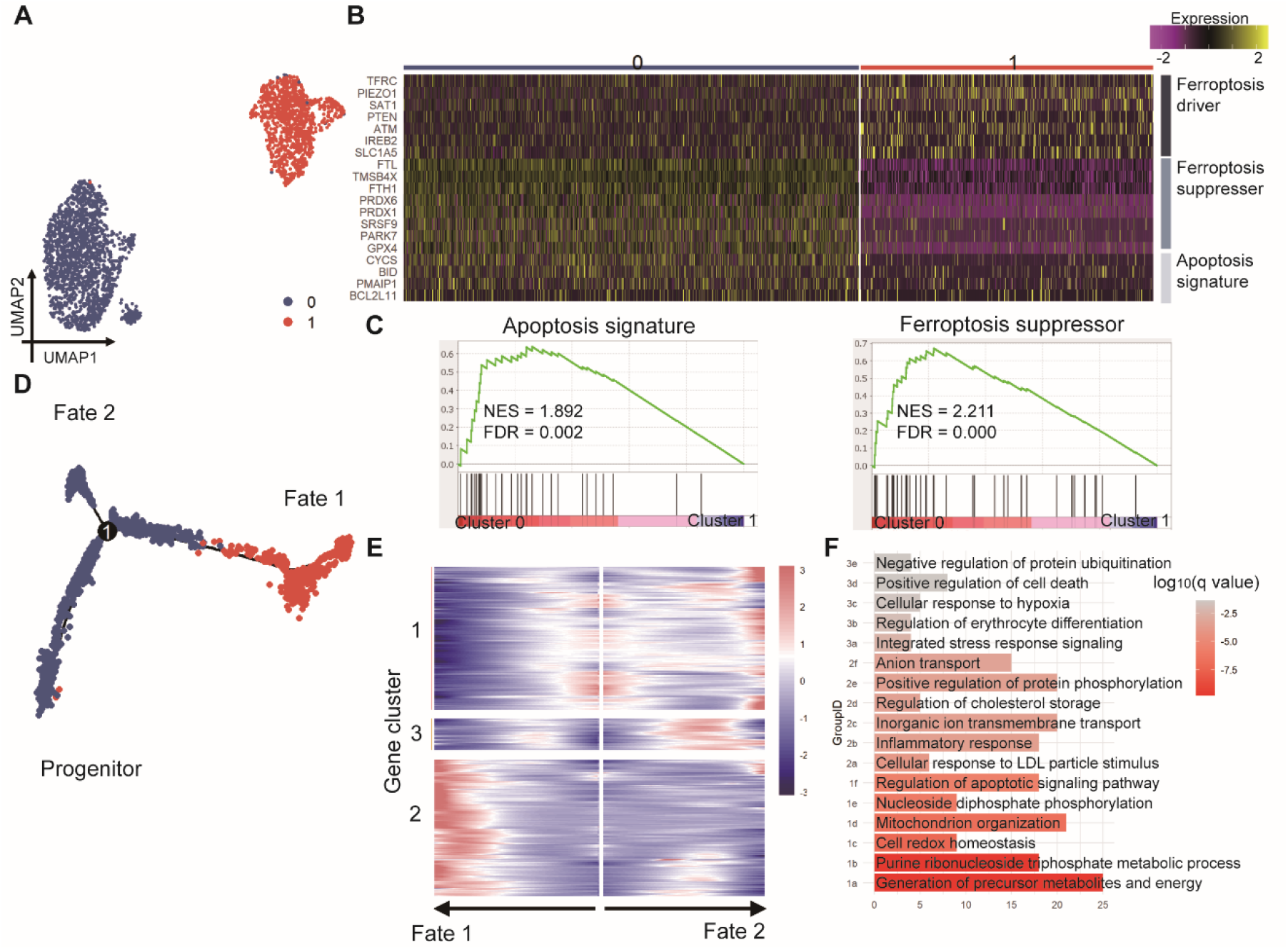
Single-cell transcriptomics depicts divergent foam cell fate. (*A*) UMAP embedding analysis of late-stage foam cells colored by the assigned clusters. (*B*) Heatmap of ferroptotic and apoptotic marker genes. (*C*) GSEA plots of apoptosis signature and ferroptotic suppressor genes. (*D*) Trajectory analysis of late-stage foam cells colored by the assigned clusters. (*E*) Heatmap showing branched gene expression based on the trajectory. (*F*) Enrichment analysis of the gene clusters defined in (*E*). TFRC, Transferrin Receptor; PIEZO1, Piezo Type Mechanosensitive Ion Channel Component 1; SAT1, Spermidine/Spermine N1-Acetyltransferase 1; PTEN, Phosphatase and tensin homolog; ATM, ataxia telangiectasia-mutated gene; IREB2, Iron Responsive Element Binding Protein 2; SLC1A5, solute carrier family 1 member 5; FTL, Ferritin Light Chain; TMSB4X, Thymosin Beta 4 X-Linked; FTH1, Ferritin Heavy Chain 1; PRDX6, Peroxiredoxin 6; PRDX1, Peroxiredoxin 1; SRSF9, Serine And Arginine Rich Splicing Factor 9; PARK7, Parkinsonism Associated Deglycase; CYCS, Cytochrome C, Somatic; BID, BH3 Interacting Domain Death Agonist; PMAIP1, Phorbol-12-myristate-13-acetate-induced protein 1; BCL2L11, Bcl-2-like protein 11.

Trajectory analysis identifies a flow toward two branches of ferroptosis and apoptosis (Fig. 6*D*). Branch analysis captured gene waves departing for the two ends and revealed three gene clusters (Fig. 6*E*). Then, enrichment analysis of Gene Ontology Biological Processes (GOBP) of the gene clusters was applied, delineating biological processes including redox, inflammation, lipid uptake and cell death (Fig. 6*F*). For gene cluster 1 (n = 256), genes involved in cell redox homeostasis (GroupID: 1c) showed a maintained down-regulation along the route to ferroptosis while genes involved in regulation of apoptotic signaling pathways (1f) were up-regulated along the route to apoptosis. The branched wave resonated with the divergent cell fate based on ferroptotic and apoptotic markers. The enrichment of purine metabolism (1b, 1e) in the branched wave suggested the potential regulatory role of purinergic signaling in a hypoxia-dependent or inflammation-dependent manner^52^. The enrichment of mitochondrion organization (1d) confirmed the central role of mitochondrial composition, metabolism and interaction with other organelles in both ferroptosis and apoptosis^42,53^. For gene cluster 2 (n = 244), genes involved in inflammatory response (2b), ion transport (2c, 2f) and lipid uptake (2a, 2d) were up-regulated along the route to ferroptosis. The enrichment of inflammatory response resonated with the crosstalk between inflammation and ferroptosis. The enrichment of ion transport suggested the plasma membrane damage in ferroptosis^54^. For gene cluster 3 (n = 57), genes involved in cellular response to hypoxia (3c) experienced an early oscillation in the way to apoptosis, suggesting that hypoxia may act as a rheostat-like regulator between ferroptosis and apoptosis^55,56^. By clustering and trajectory analysis, we verified the divergent cell fate residing in late-stage foam cells and recognized the gene regulation during the fate decision.

## Discussion

Our study integrates dynamic metabolomic and lipidomic profiling to study the metabolic and lipid reprogramming of macrophages during foam cell formation. Although previous studies recognized that oxLDL-induced foam cells acquire a M1-like phenotype^37^, we dissected the metabolic dysregulation differed from the classical inflammatory M1 phenotype and identified the dual regulation of inflammation and ferroptosis. We further integrated single-cell metabolite, lipid and RNA analysis to explore the heterogeneity during foam cell formation.

In this study, we used the THP-1 derived macrophages and foam cells to mimic the foam cell formation in atherosclerosis. Although *in vitro* models are often considered oversimplified for the high homogeneity of immortalized cell lines and a loss of heterogeneity related to microenvironment or cell-to-cell interactions^57^, our study demonstrated the heterogeneity of THP-1 derived foam cells upon exposure to identical treatment. Our time-resolved single-cell analysis witnessed a rise in the variability of apoptosis and ferroptosis in the middle and late stage of foam cell formation as well as a divergent cell fate to apoptosis or ferroptosis in the late-stage foam cells, with a reciprocal loss or gain of identified apoptotic or ferroptotic markers. These observations suggest that the decision-making whereby individual cells route to alternative types of programmed death might be a response to external stimuli based on their internal state.

By analyzing the recently published single-cell transcriptomic data of human atherosclerotic plaques, the heterogeneity of apoptosis and ferroptosis were also revealed in human plaque macrophages. In the context of atherosclerosis progression, foam cell apoptosis alleviates the plaque formation by decreasing lesion cellularity in the early lesion but promotes the plaque formation by increasing plaque necrosis in the advanced lesion; foam cell ferroptosis play a pro-atherogenic role by accelerating inflammation and oxidative stress^9^. Therefore, the divergent cell fate to apoptosis and ferroptosis provide insights into switching the fate of foamy macrophages as a potential target for atherosclerosis.

In conclusion, by exploiting the single-cell multi-omic profiling, we uncovered the cell-to-cell heterogeneity in apoptosis and ferroptosis, and depicted the molecular choreography behind the divergent cell fate of foamy macrophages. Our study inspires new understandings on the programmed death of foamy macrophages in atherosclerosis. Our approach also serves as a robust tool for investigations into disease mechanisms and drug targets. We envision that the combination of knockout and overexpression experiments (KO/OE) with single-cell multi-omics would help focus on certain drug targets and establish the molecular mechanism behind the divergent cell fate to apoptosis and ferroptosis.

## Acknowledgments

This work was supported by National Natural Science Foundation of China (NSFC, 22022401, 22074022, 21934001), Chinesisch-Deutsche Zentrum für Wissenschaftsförderung (M -0614), and the Ministry of Science and Technology of China (MOST, 2020YFF0304502, 2022YFC2704300).

## Author Contributions

Y.W. did all the experiments and data analysis. Y.W. wrote the first draft of the paper. L.L. and L.Q. revised the paper. L.L. and L.Q supervised all aspects of the work. L.L. and L.Q were involved in the design of this work.

## References

1. Tabas, I. Macrophage death and defective inflammation resolution in atherosclerosis. Nat Rev Immunol 10, 36–46 (2010).

2. Moore, K. J., Sheedy, F. J. & Fisher, E. A. Macrophages in atherosclerosis: a dynamic balance. Nat Rev Immunol 13, 709–721 (2013).

3. Moore, K. J. & Tabas, I. Macrophages in the Pathogenesis of Atherosclerosis. Cell 145, 341–355 (2011).

4. Libby, P., Ridker, P. M. & Hansson, G. K. Progress and challenges in translating the biology of atherosclerosis. Nature 473, 317–325 (2011).

5. Duewell, P. et al. NLRP3 inflammasomes are required for atherogenesis and activated by cholesterol crystals. Nature 464, 1357–1361 (2010).

6. Boyle, J. J. et al. Activating Transcription Factor 1 Directs Mhem Atheroprotective Macrophages Through Coordinated Iron Handling and Foam Cell Protection. Circ Res 110, 20–33 (2012).

7. Saeed, O. et al. Pharmacological Suppression of Hepcidin Increases Macrophage Cholesterol Efflux and Reduces Foam Cell Formation and Atherosclerosis. ATVB 32, 299–307 (2012).

8. Liu, X., Tang, Y., Cui, Y., Zhang, H. & Zhang, D. Autophagy is associated with cell fate in the process of macrophage-derived foam cells formation and progress. J Biomed Sci 23, 57 (2016).

9. Liu, C., Jiang, Z., Pan, Z. & Yang, L. The Function, Regulation and Mechanism of Programmed Cell Death of Macrophages in Atherosclerosis. Front. Cell Dev. Biol. 9, 516–527 (2022).

10. Zenobi, R. Single-Cell Metabolomics: Analytical and Biological Perspectives. Science 342, 1243259–1243259 (2013).

11. Green, D. R. The Coming Decade of Cell Death Research: Five Riddles. Cell 177, 1094–1107 (2019).

12. Hawkins, E. D., Markham, J. F., McGuinness, L. P. & Hodgkin, P. D. A single-cell pedigree analysis of alternative stochastic lymphocyte fates. PNAS 106, 13457–62 (2009).

13. Spencer, S. L., Gaudet, S., Albeck, J. G., Burke, J. M. & Sorger, P. K. Non-genetic origins of cell-to-cell variability in TRAIL-induced apoptosis. Nature 459, 428–432 (2009).

14. Paek, A. L., Liu, J. C., Loewer, A., Forrester, W. C. & Lahav, G. Cell-to-Cell Variation in p53 Dynamics Leads to Fractional Killing. Cell 165, 631–642 (2016).

15. Robbins, C. S. et al. Extramedullary Hematopoiesis Generates Ly-6Chigh Monocytes That Infiltrate Atherosclerotic Lesions. Circulation 125, 364–374 (2012).

16. Rahman, K. et al. Inflammatory Ly6Chi monocytes and their conversion to M2 macrophages drive atherosclerosis regression. The Journal of Clinical Investigation 127, 2904–2915 (2017).

17. Lin, J.-D. et al. Single-cell analysis of fate-mapped macrophages reveals heterogeneity, including stem-like properties, during atherosclerosis progression and regression. JCI Insight 4, e124574–e124590 (2019).

18. Örd, T. et al. Single-Cell Epigenomics and Functional Fine-Mapping of Atherosclerosis GWAS Loci. Circ Res 129, 240–258 (2021).

19. Winkels, H. et al. Atlas of the Immune Cell Repertoire in Mouse Atherosclerosis Defined by Single-Cell RNA-Sequencing and Mass Cytometry. Circ Res 122, 1675–1688 (2018).

20. Fernandez, D. M. Single-cell immune landscape of human atherosclerotic plaques. Nature Medicine 25, 1576– 1588 (2019).

21. Koelwyn, G. J., Corr, E. M., Erbay, E. & Moore, K. J. Regulation of macrophage immunometabolism in atherosclerosis. Nat Immunol 19, 526–537 (2018).

22. Remmerie, A. & Scott, C. L. Macrophages and lipid metabolism. Cellular Immunology 330, 27–42 (2018).

23. Haschemi, A. et al. The Sedoheptulose Kinase CARKL Directs Macrophage Polarization through Control of Glucose Metabolism. Cell Metabolism 15, 813–826 (2012).

24. Jha, A. K. et al. Network Integration of Parallel Metabolic and Transcriptional Data Reveals Metabolic Modules that Regulate Macrophage Polarization. Immunity 42, 419–430 (2015).

25. Merlin, J. et al. Non-canonical glutamine transamination sustains efferocytosis by coupling redox buffering to oxidative phosphorylation. Nat Metab 3, 1313–1326 (2021).

26. Argus, J. P. et al. Development and Application of FASA, a Model for Quantifying Fatty Acid Metabolism Using Stable Isotope Labeling. Cell Reports 25, 2919–2934.e8 (2018).

27. Dennis, E. A. et al. A Mouse Macrophage Lipidome. Journal of Biological Chemistry 285, 39976–39985 (2010).

28. Yu, X.-H. Foam cells in atherosclerosis. Clinica Chimica Acta 424, 245–252 (2013).

29. Halbrook, C. J. et al. Macrophage-Released Pyrimidines Inhibit Gemcitabine Therapy in Pancreatic Cancer. Cell Metabolism 29, 1390–1399.e6 (2019).

30. Holthuis, J. C. M. & Menon, A. K. Lipid landscapes and pipelines in membrane homeostasis. Nature 510, 48–57 (2014).

31. Hirama, T. et al. Membrane curvature induced by proximity of anionic phospholipids can initiate endocytosis. Nat Commun 8, 1393–1407 (2017).

32. Maekawa, M. & Fairn, G. D. Complementary probes reveal that phosphatidylserine is required for the proper transbilayer distribution of cholesterol. Journal of Cell Science 128, 1422–1433 (2015).

33. Lemke, G. How macrophages deal with death. Nature Reviews Immunology 19, 539–549 (2019).

34. Puchalska, P. et al. Isotope Tracing Untargeted Metabolomics Reveals Macrophage Polarization-State-Specific Metabolic Coordination across Intracellular Compartments. iScience 9, 298–313 (2018).

35. Grajeda-Iglesias, C. & Aviram, M. Specific Amino Acids Affect Cardiovascular Diseases and Atherogenesis via Protection against Macrophage Foam Cell Formation: Review Article. Rambam Maimonides Med J 9, e0022 (2018).

36. Bories, G. F. P. & Leitinger, N. Macrophage metabolism in atherosclerosis. FEBS Lett 591, 3042–3060 (2017).

37. Chinetti-Gbaguidi, G. Macrophage subsets in atherosclerosis. Nature Reviews Cardiology 12, 10–17 (2015).

38. Wang, H. Aspartate Metabolism Facilitates IL-1β Production in Inflammatory Macrophages. Frontiers in Immunology 12, 13 (2021).

39. Rath, M., Muller, I., Kropf, P., Closs, E. I. & Munder, M. Metabolism via Arginase or Nitric Oxide Synthase: Two Competing Arginine Pathways in Macrophages. Front. Immunol. 5, 532–537 (2014).

40. Palmieri, E. M. et al. Nitric oxide orchestrates metabolic rewiring in M1 macrophages by targeting aconitase 2 and pyruvate dehydrogenase. Nat Commun 11, 698–714 (2020).

41. Kapralov, A. A. et al. Redox lipid reprogramming commands susceptibility of macrophages and microglia to ferroptotic death. Nat Chem Biol 16, 278–290 (2020).

42. Gao, M. et al. Role of Mitochondria in Ferroptosis. Molecular Cell 73, 354–363.e3 (2019).

43. Braverman, N. E. & Moser, A. B. Functions of plasmalogen lipids in health and disease. Biochimica et Biophysica Acta 9, 1442–1552 (2012).

44. Zhang, D., Qin, Q. & Qiao, L. Mass spectrometry profiling of single bacterial cells reveals metabolic regulation during antibiotics induced bacterial filamentation. 8 (2022).

45. Qiao, L. et al. Electrostatic-Spray Ionization Mass Spectrometry. Anal. Chem. 84, 7422–7430 (2012).

46. Vance, J. E. & Tasseva, G. Formation and function of phosphatidylserine and phosphatidylethanolamine in mammalian cells. Biochimica et Biophysica Acta (BBA) -Molecular and Cell Biology of Lipids 1831, 543–554 (2013).

47. Zhang, L., Richard, A. S., Jackson, C. B., Ojha, A. & Choe, H. Phosphatidylethanolamine and Phosphatidylserine Synergize To Enhance GAS6/AXL-Mediated Virus Infection and Efferocytosis. J Virol 95, e02079–20 (2020).

48. Zou, Y. et al. Plasticity of ether lipids promotes ferroptosis susceptibility and evasion. Nature 585, 603–608 (2020).

49. Balgoma, D. & Hedeland, M. Etherglycerophospholipids and ferroptosis: structure, regulation, and location. Trends in Endocrinology & Metabolism 32, 960–962 (2021).

50. Minhas, P. S. et al. Macrophage de novo NAD+ synthesis specifies immune function in aging and inflammation. Nat Immunol 20, 50–63 (2019).

51. Depuydt, M. A. C. et al. Microanatomy of the Human Atherosclerotic Plaque by Single-Cell Transcriptomics. Circulation Research. 127, 1437–1455 (2020).

52. Burnstock, G. Purinergic Signaling in the Cardiovascular System. Circulation Research 120, 207–228 (2017).

53. Pena-Blanco, A. & Garc, A. J. Bax, Bak and beyond — mitochondrial performance in apoptosis. The FEBS Journal 285, 416–431 (2018).

54. Pedrera, L. Ferroptotic pores induce Ca2+ fluxes and ESCRT-III activation to modulate cell death kinetics. Cell Death & Differentiation 28, 1644–1657 (2021).

55. Greijer, A. E. The role of hypoxia inducible factor 1 (HIF-1) in hypoxia induced apoptosis. Journal of Clinical Pathology 57, 1009–1014 (2004).

56. Chen, X. Broadening horizons: the role of ferroptosis in cancer. ClInIcAl Oncology 18, 280–296 (2021).

57. Qin, Z. The use of THP-1 cells as a model for mimicking the function and regulation of monocytes and macrophages in the vasculature. Atherosclerosis 221, 2–11 (2012).

