## Supplementary information for "Single-cell time-resolved multi-omics reveal apoptotic and ferroptotic heterogeneity during foam cell formation"

**This PDF file includes:**

Figures S1 to S14
Table S1

**Supporting Information Text**

**Table S1. Abbreviations of lipid subclasses**

| Abbreviation | Full name | LipidMapsID |
|---|---|---|
| PC | Diacylglycerophosphocholines | GP0101 |
| PC-O | l-alkyl,2-acylglycerophosphocholines | GP0102 |
| LPC | Monoacylglycerophosphocholines | GP0105 |
| LPC-O | Monoalkylglycerophosphocholines | GP0106 |
| PE | Diacylglycerophosphoethanolamines | GP0201 |
| PE-P | l-(lZ-alkenyl),2-acylglycerophosphoethanolamines | GP0203 |
| LPE | Monoacylglycerophosphoethanolamines | GP0205 |
| LPE-P | Monoalkylglycerophosphoethanolamines | GP0206 |
| PS | Diacylglycerophosphoserines | GP0301 |
| LPS | Monoacylglycerophosphoserines | GP0305 |
| PG | Diacylglycerophosphoglycerols | GP0401 |
| LPG | Monoacylglycerophosphoglycerols | GP0405 |
| PI | Diacylglycerophosphoinositols | GP0601 |
| LPI | Monoacylglycerophosphoinositols | GP0605 |
| Sph | Sphing-4-enines (Sphingosines) | SP0101 |
| Cer | N-acylsphinsosines (ceramides) | SP0201 |
| GlcCer/LacCer | Simple Glc series | SP0501 |
| SM | Ceramide phosphocholines (sphingomyelins) | SP0301 |
| ST | Sterols | ST01 |
| FA | Fatty acids and conjugates | FA01 |
| TG | Triacylglycerols | GL0301 |
| DG | Diacylglycerols | GL0201 |

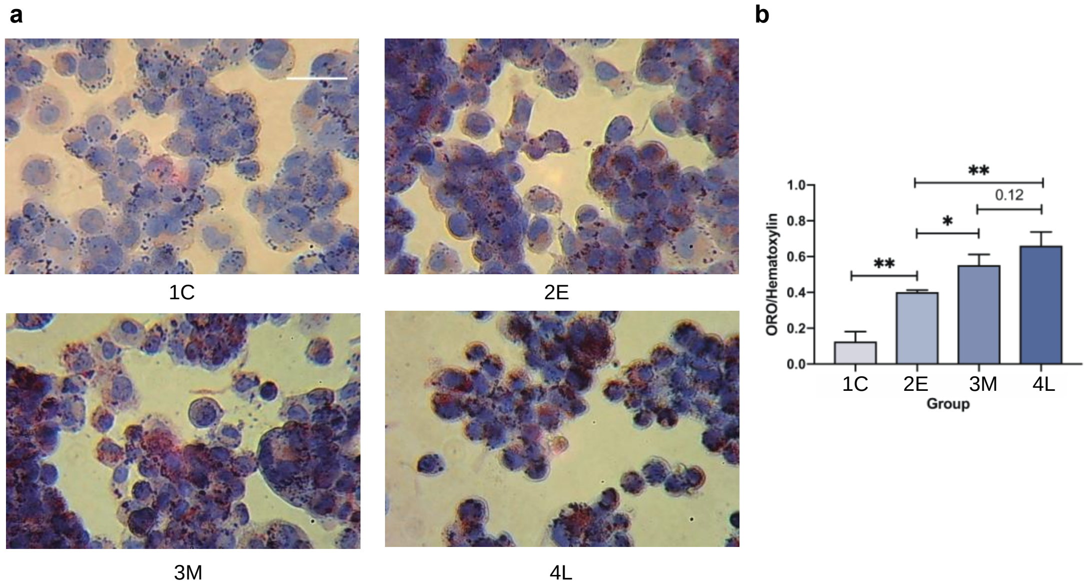


**Fig. S1. Verification of foam cell formation.** A) Morphology of the macrophages (1C) and foam cells of three stages (2E, 3M and 4L) after oil red O and hematoxylin stain. The scale bar represents 50 μm. B) Quantitative evaluation of the ORO stain. Error bars represent the standard deviation, and "*" and "**" represent p values less than 0.05 and 0.01, respectively.

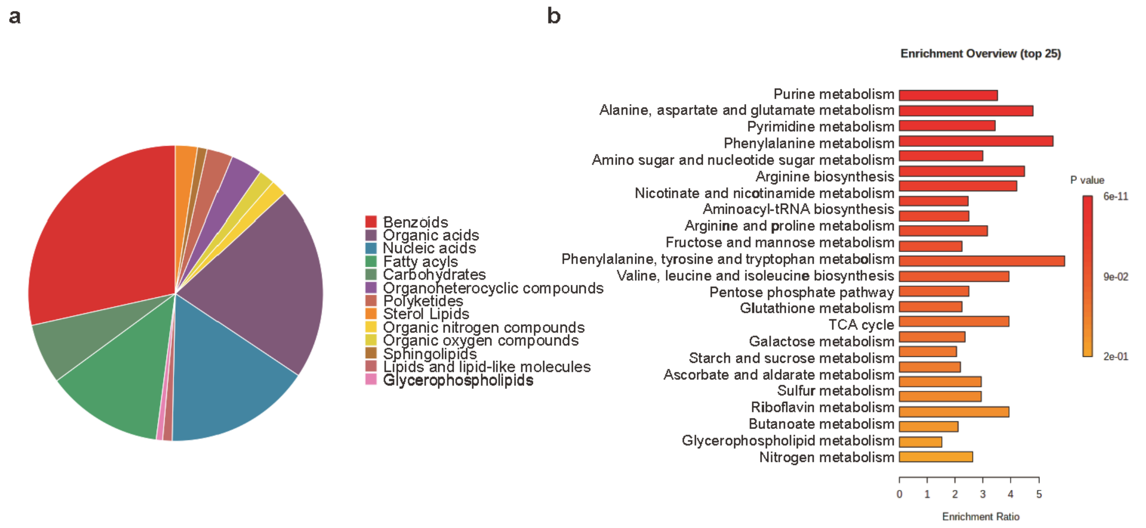


**Fig. S2. Enrichment analysis of metabolites identified from population of cells.** A) Pie chart of the superclass enrichment result. B) Bar plot of the pathway enrichment result.

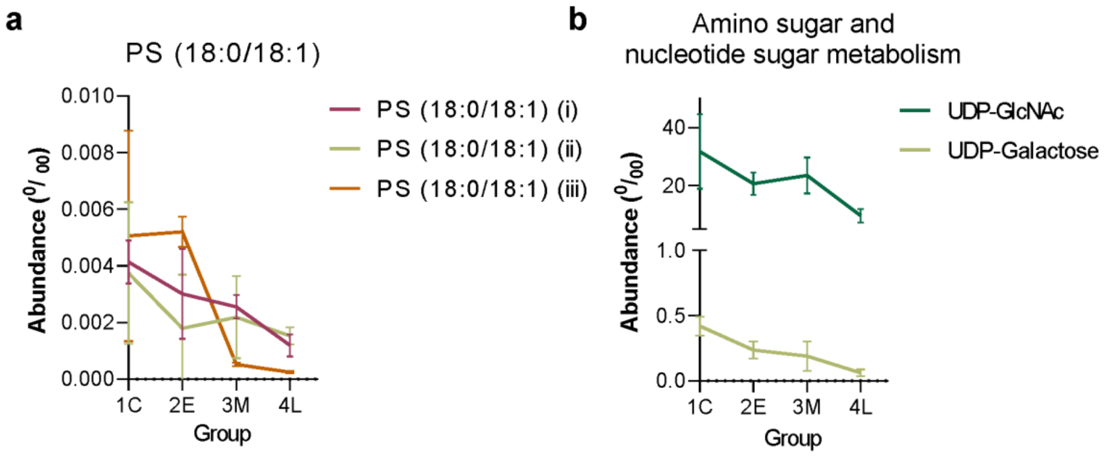


**Fig. S3. Dynamic changes of specific lipid molecular species and metabolites.** A) Dynamic changes in PS (18:0/18:1). i, ii, iii represent lipid positional and geometric isomers of C=C bonds. B) Dynamic changes of uridine diphosphate N-acetylglucosamine (UDP-GlcNAc) and uridine diphosphate galactose (UDP-Galactose) involving in amino sugar and nucleotide sugar metabolism. Error bars represent standard deviation.

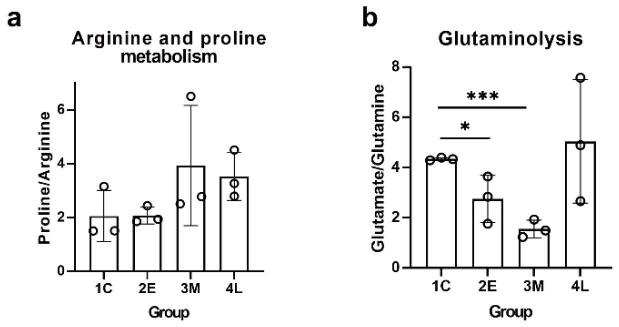


**Fig. S4. Dynamic changes in amino acid anaplerosis.** A) Dynamic changes in Proline/Arginine ratio. B) Dynamic changes in Glutamate/Glutamine ratio. Error bars represent standard deviation. "*" and "***" represent p value less than 0.05 and 0.001, respectively.

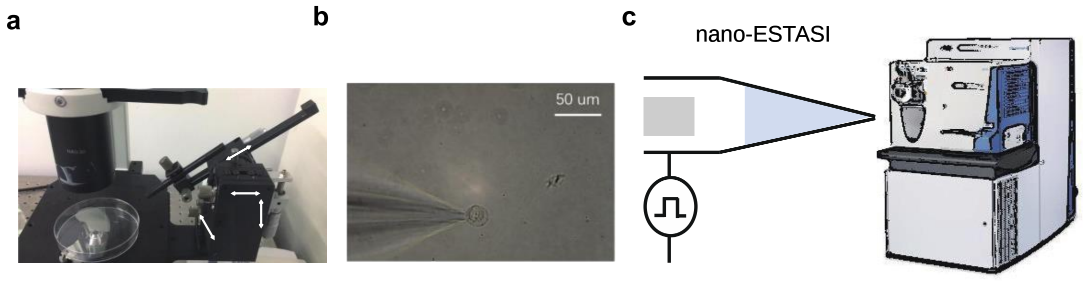


**Fig. S5. Setup for single cell MS analysis.** A) Photo of the micromanipulation stage. Arrows mark the direction of 4 axes. B) Live single cell sampling. The scale bar represents 50 µm. C) Scheme of nano-electrostatic spray ionization coupled to a mass spectrometer.

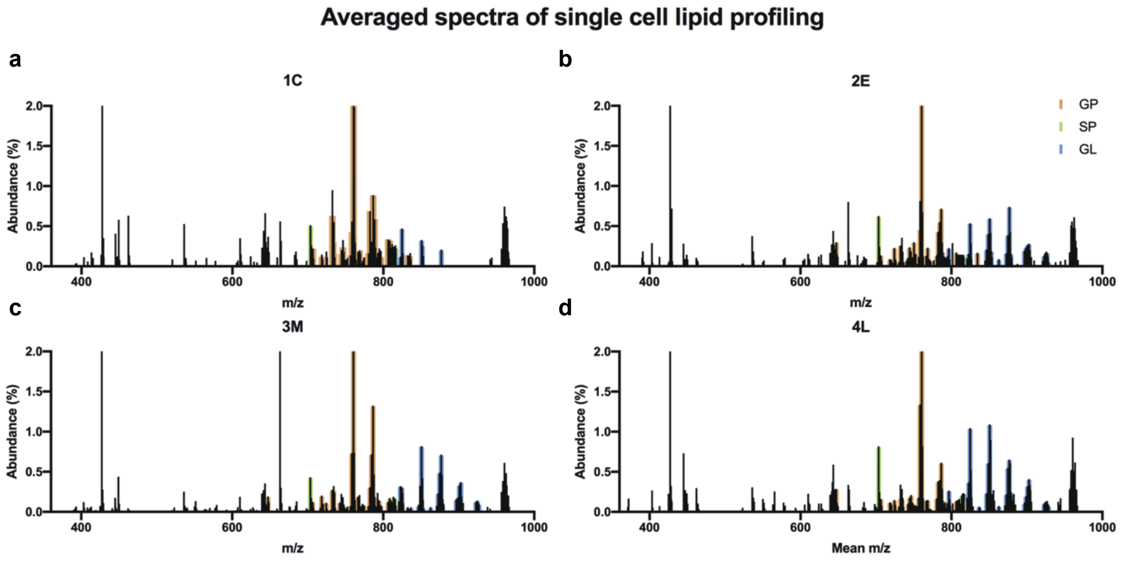


**Fig. S6. Single cell lipid profiling of macrophages and foam cells.** Annotated lipid species are labelled in color according to their categories. GP, glycerophospholipids; SP, sphingolipids; GL, glycerolipids.

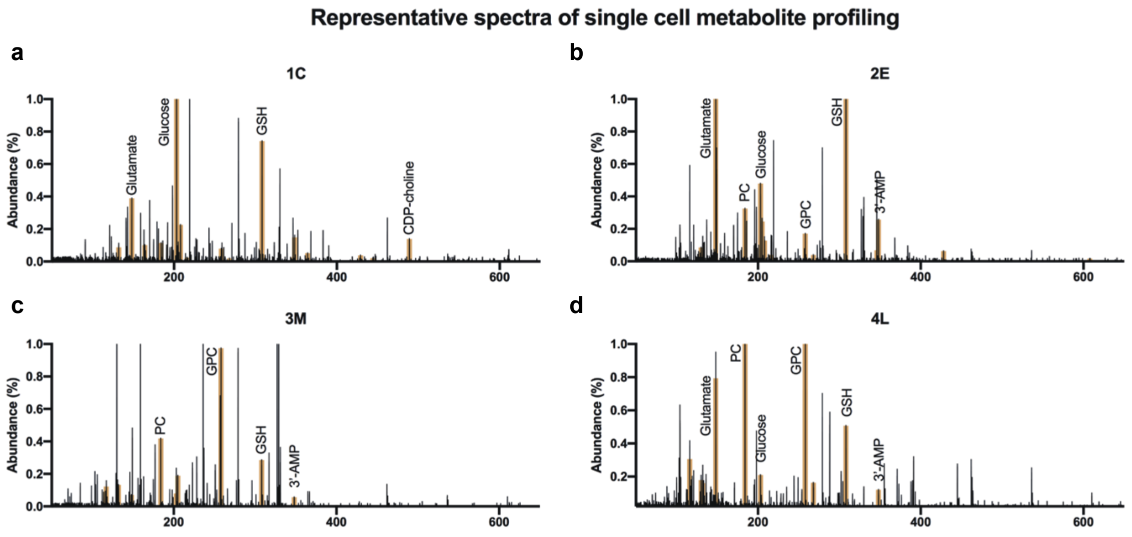


**Fig. S7. Single cell metabolite profiling of macrophages and foam cells.** Some abundant metabolites are annotated in the spectra. PC, phosphorylcholine; GPC, glycerophosphocholine; GSH, glutathione; 3’-AMP, 3’-adenosine monophosphate.

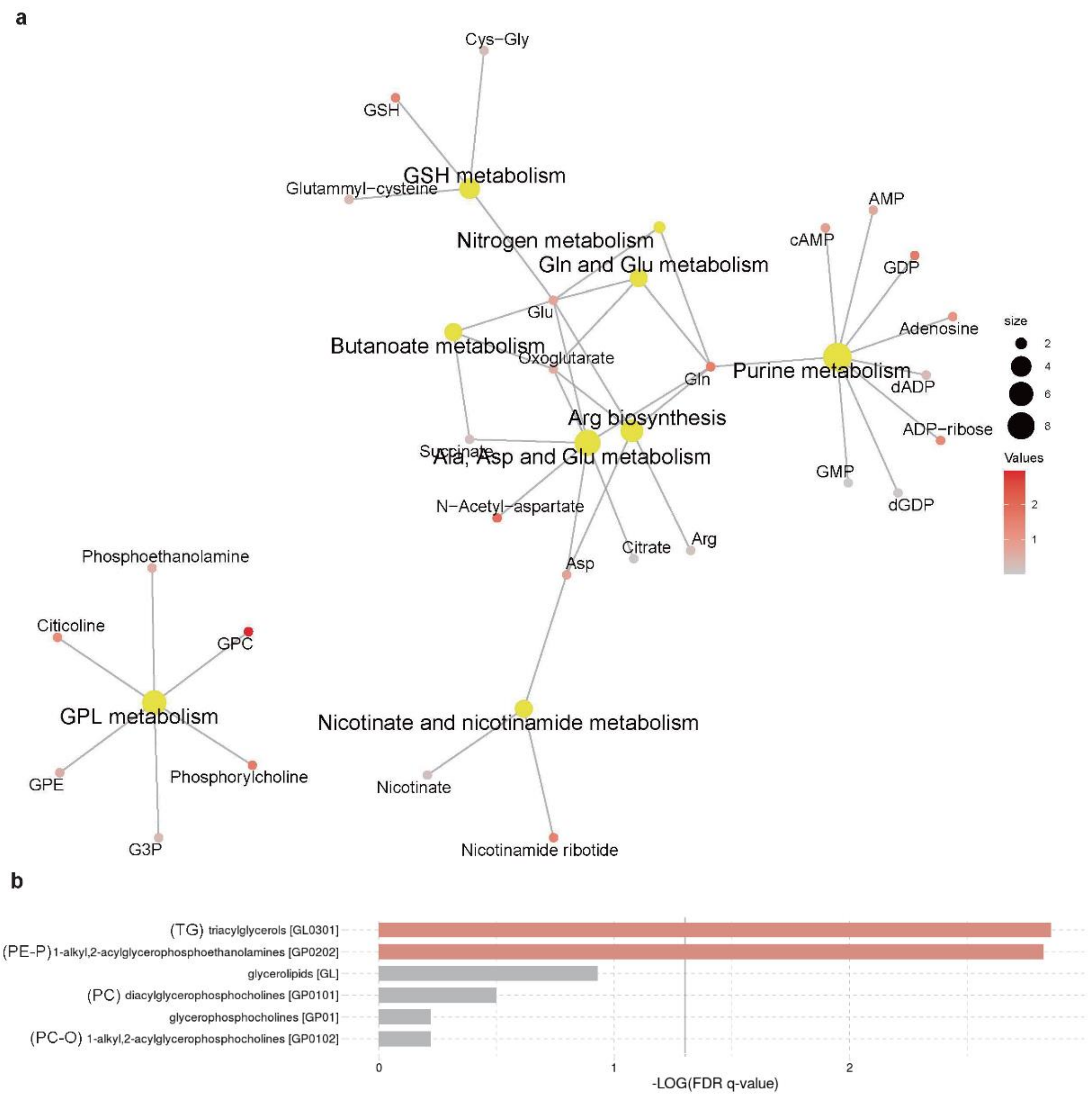


**Fig. S8. Enrichment analysis of the metabolites and lipids identified from single cells.** A) Pathway enrichment network of the single-cell metabolites. The metabolic pathways are colored in yellow and their size represent the hits in the pathway. The metabolites are colored by their VIP values based on PLSDA shown in **Fig. S10c** and **10d**. B) Subclass enrichment bar plot of the single-cell lipids. Bars colored in red suggest enrichment with q-value less than 0.05.

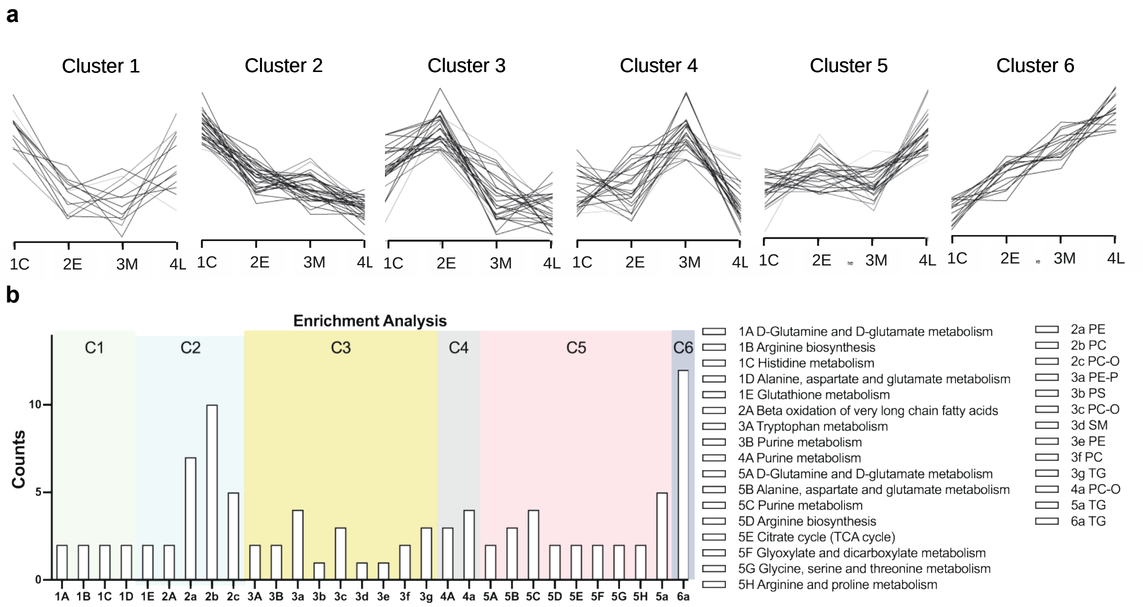


**Fig. S9. Time-series clustering analysis of the metabolites and lipids identified from single cells.** A) Clustering of the metabolites and lipids identified by single-cell MS analysis. B) Metabolic pathway and lipid sub-classes enrichment analysis of the clusters in a. 1A represents metabolites in cluster 1, while 1a represents lipids in cluster 1. The others follow the same rule.

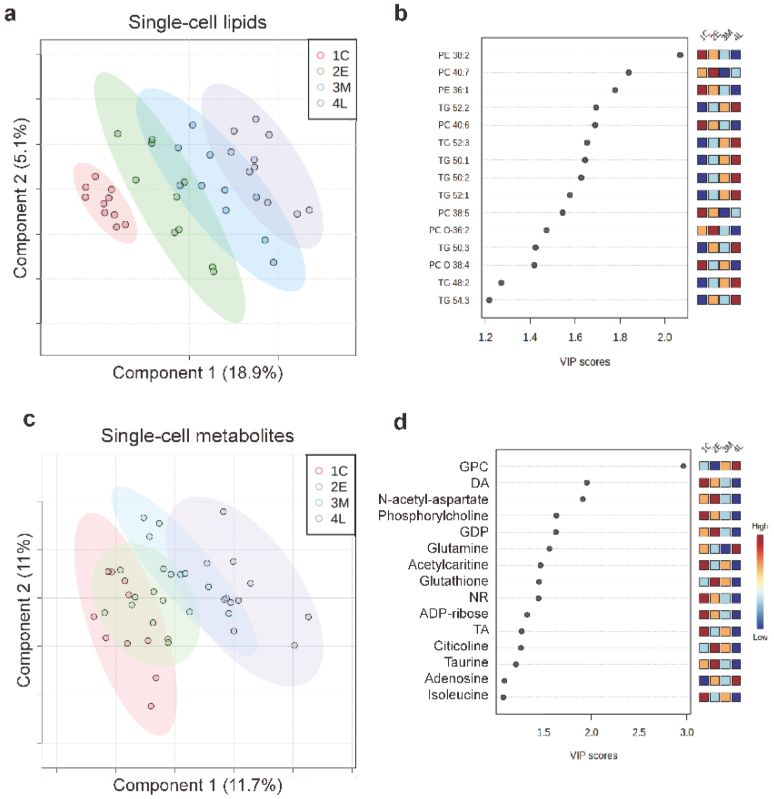


**Fig. S10. PLSDA analysis of macrophages and foam cells of three stages based on single-cell metabolites and lipid species.** A) PLSDA score plot on single-cell lipid species. B) Top 15 VIP plot of single-cell lipid species. C) PLSDA score plot on single-cell metabolites. D) Top 15 VIP plot of single-cell metabolites. GPC, glycerophsphocholine; DA, dodecanoic acid; GDP, guanosine diphosphate; NR, nictonamide ribose; ADP-ribose, adenosine diphosphate-ribose; TA, traumatic acid.

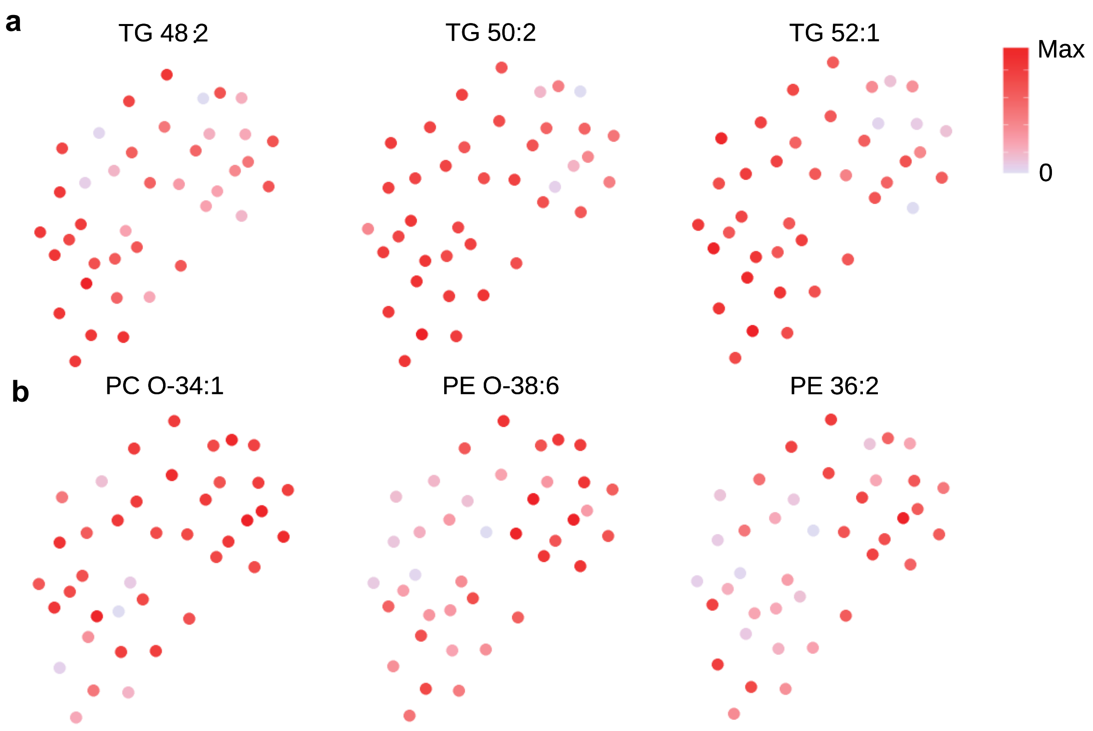


**Fig. S11. UMAP analysis on single-cell lipids.** A) UMAP layout colored by several representative TG species. B) UMAP layout colored by several ether phospholipids and PE species. The UMAP layouts were colored by the abundance of lipid features.

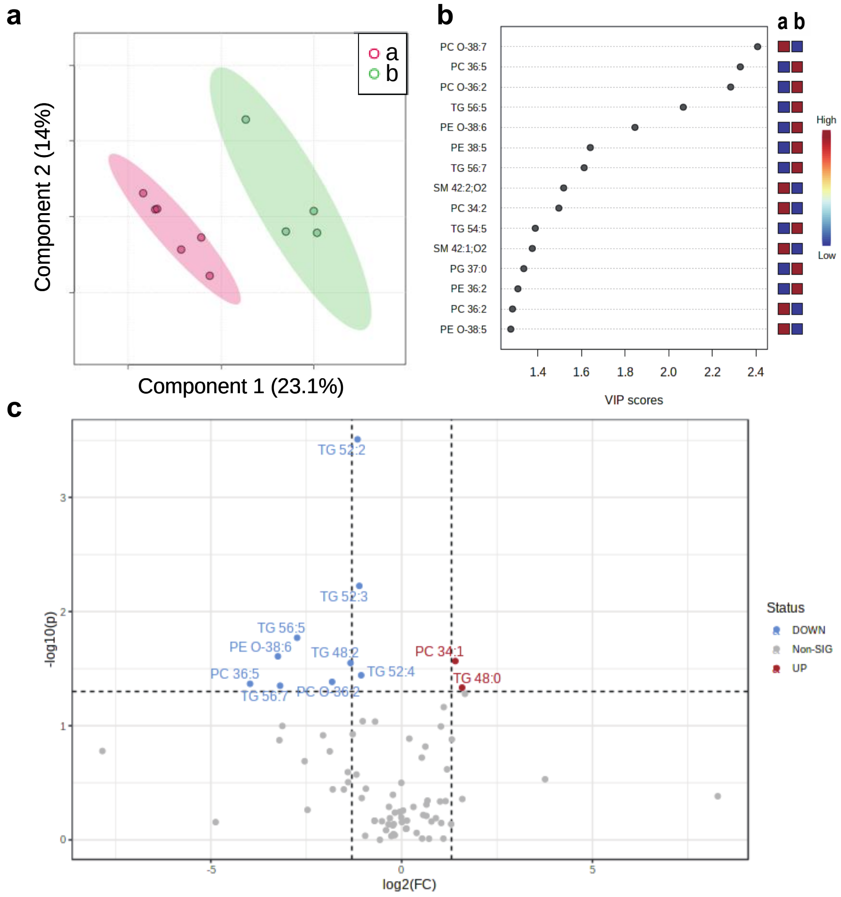


**Fig. S12. PLSDA analysis and volcano plot of the two subpopulations of late-stage foam cells as demonstrated in Fig. 4d based on single cell lipids.** A) PLSDA score plot on single-cell lipids. Group a and b represent the two subpopulations in late-stage foam cells. B) VIP plot of the single-cell lipids. c, Volcano plot between the two subpopulations based on single-cell lipids.

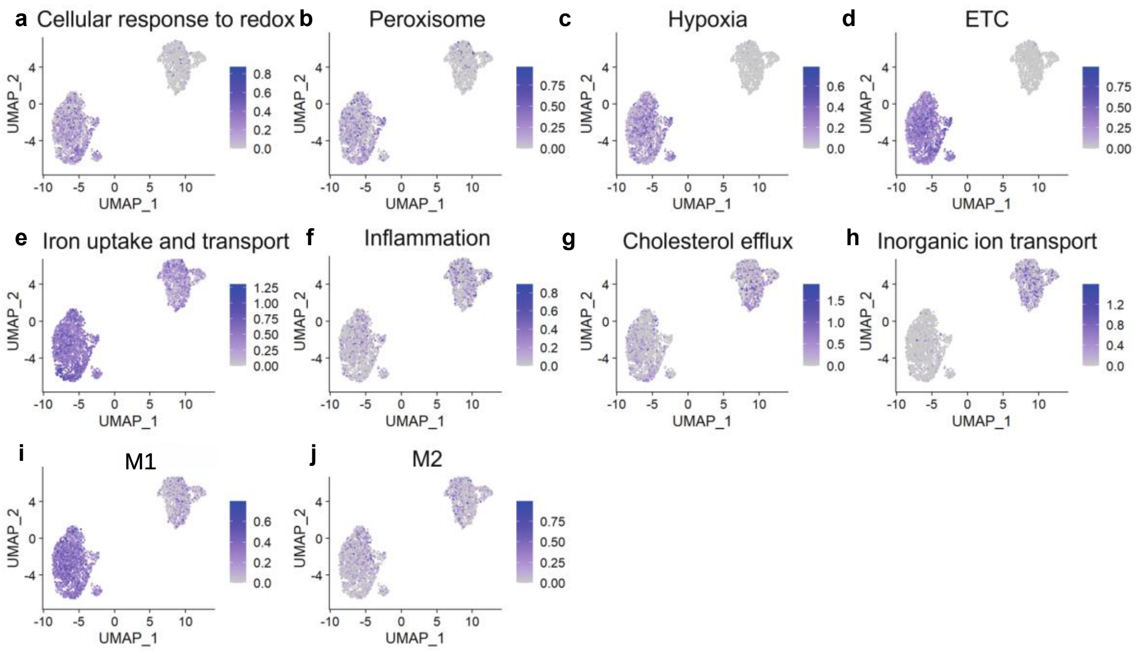


**Fig. S13. Single-cell gene set enrichment in late-stage foam cells clusters.** Single-cell transcriptomes of the two clusters were analyzed for the enrichment of specific genes and pathways. The expression of genes pertaining to cell redox homeostasis A M12813), peroxisome B M6391), cellular response to hypoxia C M641), electron transfer chain in mitochondria D M39417), iron uptake and transport E M962), acute inflammatory response F M6557), regulation of cholesterol efflux G M11164), inorganic ion import across plasma membrane H M24874), M1 signatures I M6595) and M2 signatures J M6596).

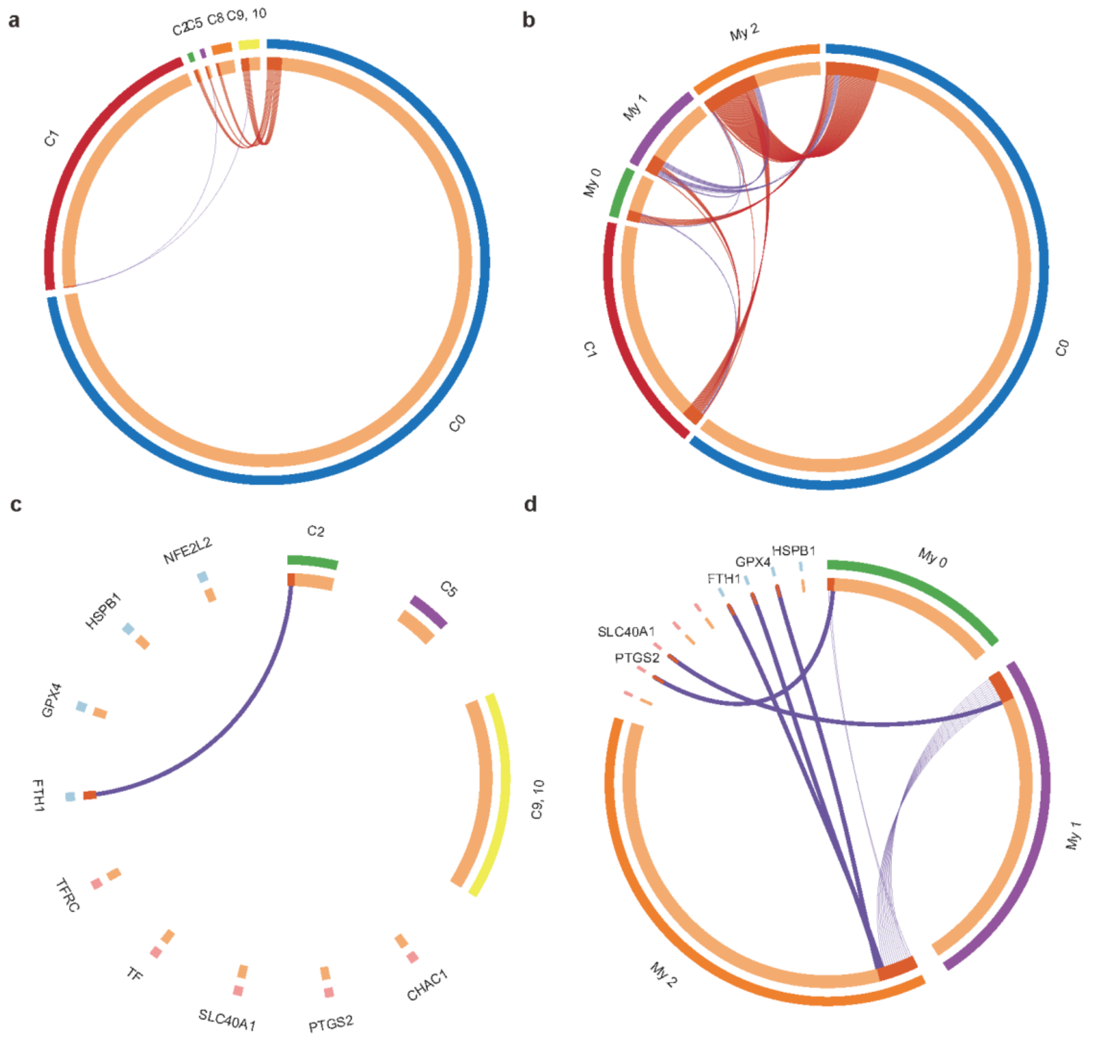


**Fig. S14. Circos plots showing the overlap of macrophage clusters between the *in vitro* dataset and clinical datasets.** A) Overlap with Fernandez et al.'s publication. C0 and C1 represent the two clusters identified in our dataset; C2, C5, C6, C8 and C9, 10 represent the macrophage clusters in Fernandez et al.'s publication. Hypergeometric test showed significant C0-C2, C0-C5 and C0-C9, 10 overlap. B) Overlap with Depuydt et al.'s publication. C0 and C1 represent the two clusters identified in our dataset; My 0, My 1 and My 2 represent the macrophage clusters in Depuydt et al.'s publication. Hypergeometric test showed significant C0-My 0, C1-My 1, C0-My 2 and C1-My 2 overlap. The significant and insignificant overlap were colored in red and purple, respectively. C0: apoptosis-like macrophages; C1: ferroptosis-like macrophages; C2: M(Hb)-like macrophages (symptomatic); C5: foamy macrophages (asymptomatic); C8, pro-inflammatory macrophages (asymptomatic); C9, 10 M2-like macrophages (symptomatic); My 0: $IL1B^{+}$ inflammatory macrophages; My 1: $TNF^{+}$ inflammatory macrophages; My2: $ABCG1^{+}$ foamy macrophages. C,D) Overlap with ferroptosis marker genes. Ferroptosis driver genes and suppressor genes were labelled in red and blue, respectively. Bands represent a hit with a marker gene, while lines represent intracluster overlap. CHAC1, ChaC glutathione specific gamma-glutamylcyclotransferase 1; PTGS2, prostaglandin-endoperoxide synthase 2; SLC40A1, solute carrier family 40 member 1; TF, Transferrin; TFRC, Transferrin Receptor; FTH1, Ferritin Heavy Chain 1; GPX4, glutathione peroxidase 4; HSPB1, heat shock protein family B (small) member 1; NFE2L2, NFE2 like bZIP transcription factor 2.
